# Performance of the Maxim and Sedia Limiting Antigen Avidity assays for population-level HIV incidence surveillance

**DOI:** 10.1101/572677

**Authors:** Joseph B. Sempa, Alex Welte, Michael P. Busch, Jake Hall, Dylan Hampton, Sheila M. Keating, Shelley N. Facente, Kara Marson, Neil Parkin, Christopher D. Pilcher, Gary Murphy, Eduard Grebe, on behalf of the Consortium for the Evaluation and Performance of HIV Incidence Assays (CEPHIA)

## Abstract

**Background:** Two manufacturers, Maxim Biomedical and Sedia Biosciences Corporation, supply US CDC-approved versions of the HIV-1 Limiting Antigen Avidity EIA (LAg assay) for detecting ‘recent’ HIV infection in cross-sectional incidence estimation. This study assesses and compares the performance of the Maxim and Sedia LAg assays for incidence surveillance.

**Methods:** We ran both assays on a panel of 2,500 well-characterized HIV-1-infected specimens, most with estimated dates of (detectable) infection. We analysed concordance of assay results, assessed reproducibility using repeat testing and estimated the critical performance characteristics of a test for recent infection—mean duration of recent infection (MDRI) and false-recent rate (FRR)—for a range of normalized optical density (ODn) recency discrimination thresholds, alone and in combination with viral load thresholds. We further defined three hypothetical surveillance scenarios and evaluated overall performance for incidence surveillance, defined as the precision of incidence estimates, by estimating context-specific performance characteristics.

**Results:** The Maxim assay produced lower ODn values than the Sedia assay on average, largely as a result of higher calibrator readings (mean calibrator OD of 0.749 vs. 0.643). Correlation of non-normalized OD readings was greater (*R*^2^ = 0.938) than those of ODn readings (*R*^2^ = 0.908), and the slope was closer to unity (1.054 vs. 0.899). Reproducibility of repeat testing of three blinded control specimens (25 replicates each) was slightly better for the Maxim assay (CV 8.9% to 14.8% vs. 13.2% to 15.0%). The MDRI of a Maxim-based algorithm at the ‘standard’ recency discrimination threshold in combination with viral load (ODn ≤1.5 & VL >1,000) was 201 days (95% CI: 180,223) and for Sedia was 171 days (95% CI: 152,191). Commensurately, the Maxim algorithm had a higher FRR in treatment-naive subjects (1.7% vs. 1.1%). We observed statistically significant differences in MDRI using the ODn alone (≤1.5) and in combination with viral load (>1,000). Under three fully-specified hypothetical surveillance scenarios (comparable to South Africa, Kenya and a concentrated epidemic), recent infection testing algorithms based on the two assays produced similar precision of incidence estimates.

**Conclusions:** Differences in ODn measurements between the Maxim and Sedia LAg assays on the same specimens largely resulted from differences in the reactivity of calibrators supplied by the manufacturers. Performance for surveillance purposes was extremely similar, although different ODn thresholds were optimal and different values of MDRI and FRR were appropriate for use in survey planning and incidence estimation.

## 1 Background

The Limiting Antigen Avidity EIA (LAg-Avidity Assay) was developed by the US Centers for Disease Control and Prevention (CDC) for detecting ‘recent’ HIV infection for the purposes of crosssectional incidence estimation [1]. Two major manufacturers supply versions of the assay: Maxim Biomedical (Bethesda, MD) and Sedia Biosciences Corporation (Portland, OR), with both manufacturers currently utilizing multisubtype HIV-1 recombinant antigen supplied by the CDC. A third manufacturer, Beijing King Hawk Pharmaceutical Co. (Beijing, PRC), has recently entered the market, but without US CDC involvement [2].

The Maxim and Sedia assays have generally been assumed to perform similarly, and users of the Maxim assay have mainly used performance metrics (mean duration of recent infection—MDRI— and false-recent rate—FRR) estimated from calibration data produced using the Sedia assay [3]. A recent comparison of the assays, based on 1,410 treatment-naïve specimens, found substantially lower normalized optical densities (attributed to differences in calibrators) and consequently a longer MDRI (at the ‘standard’ threshold of 1.5) for the Maxim assay [4]. We present the first largescale independent evaluation of the Maxim LAg assay for surveillance applications, including a comparative assessment of performance relative to the Sedia LAg assay, previously evaluated on the same large blinded specimen panel by our group [5, 6].

In order to assess real-world performance, we adopted the previously-proposed definition of optimal performance as the precision of incidence estimates obtainable using the algorithm under evaluation—implying a trade-off between MDRI and FRR [7]—in specified surveillance scenarios. FRR is inherently context-dependent, depending strongly on epidemiological factors such as the antiretroviral treatment coverage, abundance of elite controllers (spontaneous virus suppression) and distribution of times-since-infection in the surveyed population (see [8]). MDRI largely reflects the biological properties of the test (i.e. the post-infection biomarker dynamics), but is also affected by the specific screening assay used in a survey to ascertain HIV-positivity, the subtype mix in the population, etc. We therefore defined three hypothetical surveillance scenarios, based on real-world settings, and evaluated performance of the Maxim and LAg assays under the assumptions defining the scenarios.

## 2 Methods

### 2.1 The CEPHIA Evaluation Panel

The CEPHIA specimen repository houses more than 29,000 unique specimens from over 3,000 HIV-1-positive subjects. The CEPHIA Evaluation Panel (EP) consists of 2,500 plasma specimens [5, 6] that were obtained from 928 unique subjects (1–13 specimens per subject), spanning a wide range of times since infection. Most specimens are from subjects with HIV subtype B (57% of specimens), C (27%), A1 (10%), and D (5%). The panel further contains multiple blinded aliquots of 3 control specimens (25 replicates of each), with antibody reactivity characteristic of recent, intermediate, and long-standing infection, to allow evaluation of the reproducibility of assay results, and moderate numbers of specimens from ART-suppressed and naturally suppressed (elite controller) participants to assess the impact of viral suppression on FRR.

The majority of subjects contributing specimens to the panel (68%) had sufficient clinical background data to produce Estimated Dates of Detectable Infection (EDDIs), which are obtained by systematically interpreting diverse diagnostic testing histories into infection date ‘point estimates’ (EDDIs) and plausible intervals of first detectability according to the method previously described [9]. EDDIs represent the date on which a viral load assay with a 50% limit of detection (LoD) of 1 RNA copy/mL would be expected to first detect the infection, and consequently MDRI estimates are ‘anchored’ to this reference test.

The UCSF Institutional Review Board approved the CEPHIA study procedures (approval #10-02365), and all specimens were collected under IRB-approved research protocols.

### 2.2 Laboratory Procedures

The CEPHIA EP was tested with the Maxim and Sedia™ HIV-1 Limiting Antigen Avidity EIA assays (Maxim and Sedia LAg, respectively), according to their respective product inserts [10, 11]. Both assays are microtitre-based with the solid phase of the microtitre plate coated with a multisubtype recombinant HIV-1 antigen. This antigen is coated in a limiting concentration to prevent crosslinking of antibody binding, thereby making it easier to remove weakly-bound antibody. Specimen dilutions are incubated for 60 minutes and then a disassociation buffer is added for 15 minutes to remove any weakly-bound antibody. A goat anti-human, horseradish peroxidase (HRP)-conjugated IgG is added and this binds to any remaining IgG; a TMB substrate is added and a colour is generated which is proportionate to the amount of HRP. An optical density (OD) is measured for each sample and this is normalized by use of a calibrator specimen. On each plate, the calibrator is tested in triplicate, with the median of the three ODs used to normalize specimen readings, producing normalized optical density (ODn) measurements.

The procedures for both assays are essentially the same, and both manufacturers source the recombinant antigen from the CDC as part of their licensing agreement. However, other components of the assay, such as the type of plates used, the control and calibrator materials, etc., were sourced or produced by the individual manufacturers. The testing procedure for both assays requires that specimens producing an initial ‘screening’ OD of ≤2.0 be subjected triplicate ‘confirmatory’ testing. The median ODn of the triplicate results then serves as the final result [10, 11]. In the Maxim evaluation, a small number of specimens erroneously did not receive the triplicate confirmatory testing (940 out of 952 that should have received confirmatory testing did and 12 did not). A simulation investigation showed that this minor protocol deviation did not substantially affect our results. It is further recommended that specimens producing an initial ODn ≤0.4 be subjected to serological testing to confirm HIV infection. All subjects contributing specimens to the CEPHIA panel were confirmed HIV-1-positive, and this step could be omitted.

Laboratory technicians were blinded to specimen background data during testing, which for each of the assays was completed in batches over a one month period using LAg kits procured from the relevant manufacturer at the same time.

### 2.3 Statistical Analysis

We evaluated a Recent Infection Testing Algorithm (RITA) which consists of a screening assay to ascertain HIV infection followed by a single immunoassay (either Maxim LAg or Sedia LAg) as primary recency marker, and a quantitative viral load. Recent infection was defined as an ODn *below* a tunable threshold, and a quantitative viral load *above* a tunable threshold. In addition, the RITA requires the specification of a cut-off time, denoted *T*, with recent results obtained from individuals infected for longer than *T* defined as falsely recent [12]. A large number of ODn thresholds are investigated in addition to the ‘standard’ LAg threshold of ODn ≤1.5. Performance without a supplemental viral load, and viral load thresholds of 0 (meaning that no threshold was applied, but analysis is restricted to specimens that have viral load data available), 75, 400, 1,000, and 5,000c/mL were investigated. We used a value of 2 years for *T* throughout.

The performance of a test for recent infection in cross-sectional incidence estimation is reflected in two key characteristics: the MDRI and FRR. These characteristics and methods for estimating them are described in greater detail elsewhere [12, 5, 8]

Briefly, MDRI is the average time an individual spends in the ‘recent infection’ state as defined by the biomarker or set of biomarkers, having been infected for less than *T*. We estimated MDRI by fitting binomial regression models for the probability of exhibiting the recent marker as a function of time since detectable infection *t*, and integrated this function *P*_*R*_(*t*) from 0 to *T*. Confidence intervals were estimated by means of subject-level bootstrap resampling (10,000 iterations). MDRI may be sensitive to HIV-1 subtype, which affects post-infection antibody dynamics [5, 6, 8], so MDRIs were derived separately for different HIV subtypes in addition to an overall MDRI reflecting the CEPHIA evaluation panel, and ‘blended’ MDRIs (weighted averages of subtype-specific MDRIs) for each surveillance setting.

FRR (also referred to as the false-recent ratio) is simply the proportion of individuals infected for longer than *T* who nevertheless exhibit the ‘recent’ biomarker. The precision of incidence estimates are highly sensitive to FRR, and in most cases values above about 1% result in poor precision. As noted earlier, FRR depends strongly on context, since viral suppression, either as a result of antiretroviral treatment or spontaneous suppression, frequently results in partial seroreversion which leads to the production of false-recent results on serological markers. Inclusion of viral load in a RITA (i.e. viral load less than some threshold results in classification as long-term infection, irrespective of ODn result) ameliorates the impact of viral suppression. It is important to consider a range of RITAs, including multiple ODn and viral load threshold combinations. In practice, a viral load threshold of >1,000c/mL is frequently used, especially when dried blood spot (DBS) specimens are collected for recency ascertainment, which makes quantification of viral RNA at lower concentrations difficult. Naïve FRR estimates (i.e., not adated to epidemiological context), and their confidence intervals, were obtained by estimating the binomial probability that an untreated individual would produce a recent result when infected for longer than *T*

To evaluate performance for surveillance purposes, we estimated context-specific MDRI and FRR in three hypothetical (but realistic) scenarios. The three scenarios were defined as follows:

**Scenario A** (South Africa-like epidemic): 100% subtype C infection; HIV prevalence of 18.9% (SE: 1.12%); Incidence of 0.990 cases/100 person-years (SE: 0.0004); ART coverage and viral suppression rates of 56% (SE: 5.6%) and 82% (SE: 8.2%), respectively; survey sample size of 35,000. **Scenario B** (Kenya-like epidemic): 70% subtype A, 25% subtype D, and 5% subtype C; HIV prevalence of 5.4% (SE: 0.36%); Incidence of 0.146 cases/100PY (SE: 0.039); ART coverage and viral suppression rates of 64% (SE: 6.4%) and 81% (SE: 8.1%), respectively; survey sample size of 14,000. **Scenario C** (North American key population-like epidemic): 100% subtype B; HIV prevalence of 15.0% (SE: 1.00%); Incidence of 0.5 cases/100PY (SE: 0.050); ART coverage and viral suppression rates of 90.0% (SE: 9.0%) and 90.0% (SE: 9.0%); survey sample size of 5,000. In all scenarios, we assumed that we were able to classify all treated subjects as long-term using ART exposure testing. Therefore, the FRR in treated subjects (both suppressed and unsuppressed) is 0. This is very favourable to the RITA, but we relaxed this assumption in a sensitivity analysis reported in Figures A.4–A.6 in the Appendix. The screening assay was a laboratory assay with an average ‘diagnostic delay’ of 10.7 days relative to the 1c/mL reference test to which infection date estimates are anchored, as described above.

To obtain context-specific FRR estimates, denoted *ϵ*_*T*_, we estimated FRR in untreated individuals by fitting *P*_*R*_ (*t*) for all times post-infection and weighted it by the probability density function of times-since-infection amongst the untreated population ρ(*t*), the latter parameterized as a Weibull survival function whose shape and scale parameters were chosen to produce a weighting function consistent with prevalence and treatment coverage, and normalized to recent incidence. We estimated the FRR in treated individuals, *P*_*R*|*tx*_, as the binomial probability that a treated individual infected for longer than *T* tests recent. We then obtain a weighted FRR estimate as shown in equation 1 below.

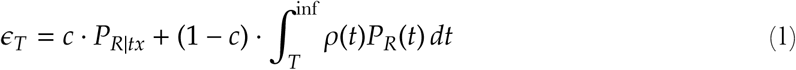

where *c* is the treatment coverage,

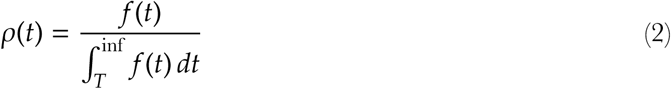

and

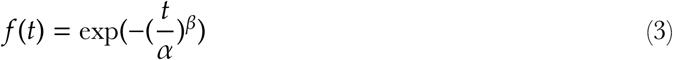

with *α* and *β* in equation 3 the Weibull scale and shape parameters, respectively. This approach was previously described in [8] and [13].

While we have declared hypothetical scenarios in which epidemiological parameters are ‘known’, we demonstrate the procedure that would be recommended in real-world settings by taking into account uncertainty in these parameters. To evaluate reproducibility of FRR estimates, we boot-strapped (30,000 iterations) both the calibration data and contextual parameters, the latter drawn from truncated normal distributions with means and standard deviations as defined for the scenarios above.

Performance was defined as the precision of incidence estimates, i.e. the relative standard error (RSE) on the incidence estimate, given the epidemiological parameters, survey sample size (assuming simple random sampling) and test characteristics (incorporating uncertainty in these), estimated for each scenario using a range of ODn thresholds and supplemental viral load thresholds, with the *inctools* R package [14] and extensions thereto [15].

## 3 Results

### 3.1 Calibrators and reproducibility on replicate control specimens

As reported in Table 1, the mean OD for all Maxim LAg calibrators was 0.75 and for Sedia LAg was 0.65, a difference of about 15.4% of the lower Sedia value. When restricted to only the calibrators used for normalization—i.e., the median value of the three ODs obtained from triplicate testing on each plate—the coefficients of variation (CVs) of Maxim and Sedia calibrators were 9.3% and 14.2%, respectively. As shown in Figure 1, Maxim calibrators produced meaningfully higher OD readings than the Sedia calibrators, with a difference in mean OD of 0.107 (95% CI: 0.090,0.123) and a p-value obtained using a Welch two-sample t-test <0.001.

**Table 1:**
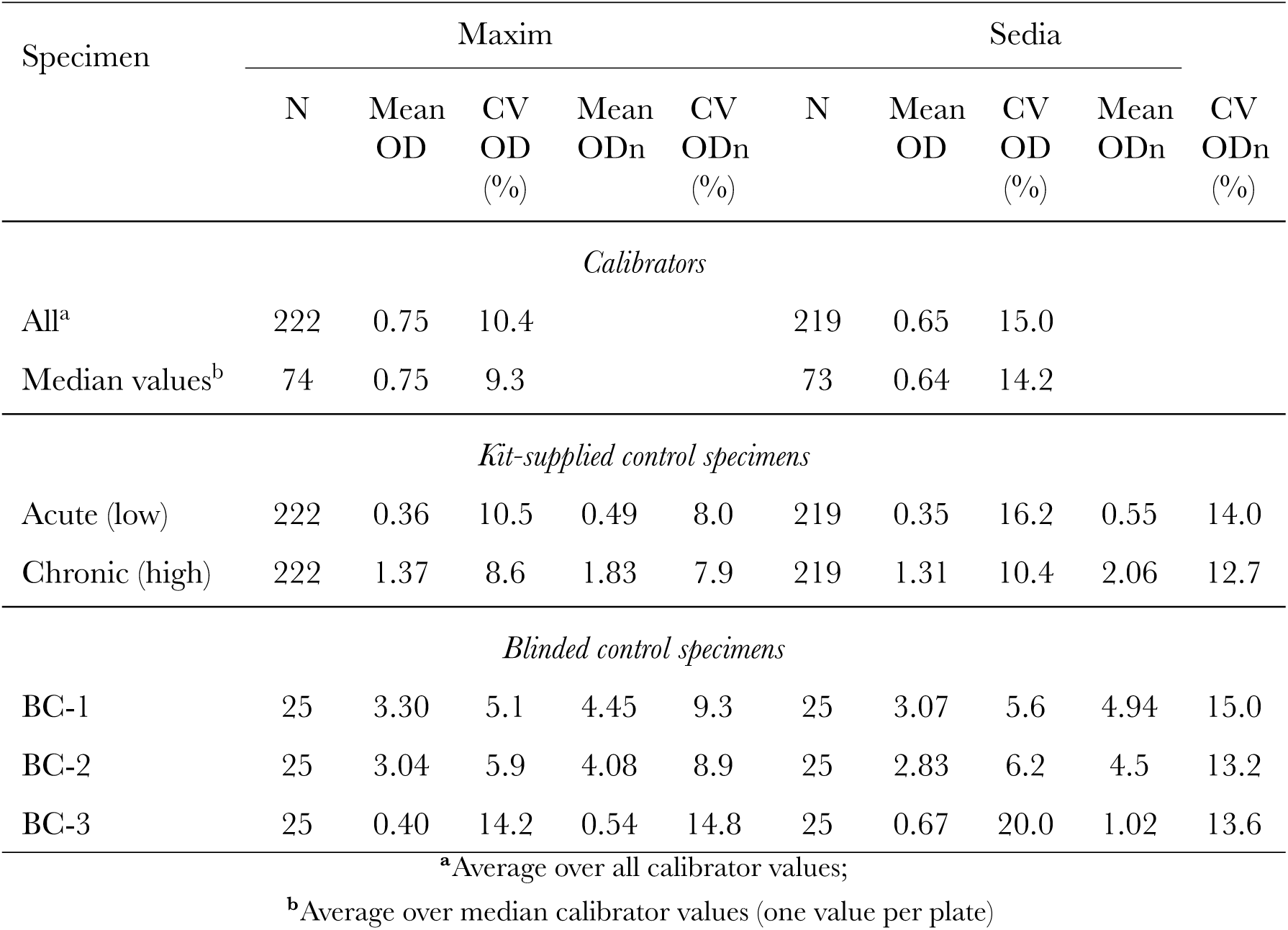
Calibrator reactivity and reproducibility of results (assessed by repeat testing)

| Specimen | Maxim |  |  |  |  | Sedia |  |  |  |  |
| --- | --- | --- | --- | --- | --- | --- | --- | --- | --- | --- |
|  | N | Mean<br>OD | CV<br>OD<br>(%) | Mean<br>ODn | CV<br>ODn<br>(%) | N | Mean<br>OD | CV<br>OD<br>(%) | Mean<br>ODn | CV<br>ODn<br>(%) |
| *Calibrators* |  |  |  |  |  |  |  |  |  |  |
| All<sup>a</sup> | 222 | 0.75 | 10.4 |  |  | 219 | 0.65 | 15.0 |  |  |
| Median values<sup>b</sup> | 74 | 0.75 | 9.3 |  |  | 73 | 0.64 | 14.2 |  |  |
| *Kit-supplied control specimens* |  |  |  |  |  |  |  |  |  |  |
| Acute (low) | 222 | 0.36 | 10.5 | 0.49 | 8.0 | 219 | 0.35 | 16.2 | 0.55 | 14.0 |
| Chronic (high) | 222 | 1.37 | 8.6 | 1.83 | 7.9 | 219 | 1.31 | 10.4 | 2.06 | 12.7 |
| *Blinded control specimens* |  |  |  |  |  |  |  |  |  |  |
| BC-1 | 25 | 3.30 | 5.1 | 4.45 | 9.3 | 25 | 3.07 | 5.6 | 4.94 | 15.0 |
| BC-2 | 25 | 3.04 | 5.9 | 4.08 | 8.9 | 25 | 2.83 | 6.2 | 4.5 | 13.2 |
| BC-3 | 25 | 0.40 | 14.2 | 0.54 | 14.8 | 25 | 0.67 | 20.0 | 1.02 | 13.6 |

<sup>a</sup>Average over all calibrator values;
<sup>b</sup>Average over median calibrator values (one value per plate)

**Figure 1:**
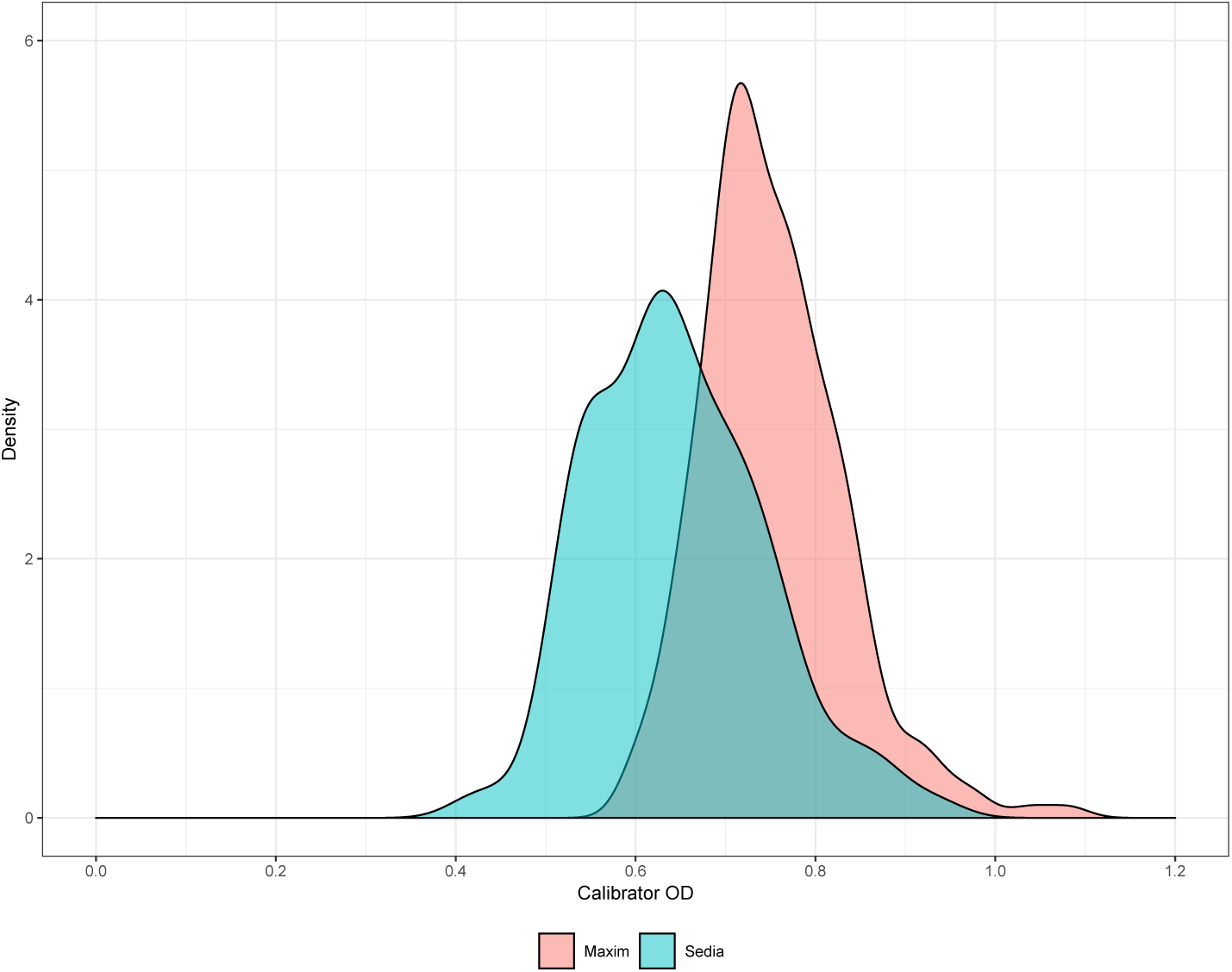
Density plot of Maxim and Sedia calibrator ODs.

Reproducibility on the three blinded control specimens was similar, with CVs on OD and ODn (across 25 replicates) slightly higher for Sedia. The Maxim assay produced lower ODn values on average, and a much lower mean ODn on the low-reactivity specimen (labelled BC-3), of 0.54 vs. 1.02 on the Sedia assay. In accordance with the manufactures’ instructions for use, specimen BC-3 was subjected to triplicate confirmatory testing on both assays. The reported ODs were those obtained from the initial screening runs, and the mean and CV on ODn results were computed on the 25 final values.

### 3.2 Performance on clinical specimens

Figure 2 shows results of testing clinical specimens in the CEPHIA EP and the impact of the previously-noted higher Maxim calibrator readings. ODn values in Figure 2B are concentrated below the diagonal line, especially at lower ODn values in the range of plausible recency discrimination thresholds. In fact, correlation was stronger for non-normalized OD readings than for normalized ODn readings. The slope for OD in Figure 2A (1.048, 95% CI: 1.037,1.059) is closer to unity than the slope for ODn in Figure 2B (0.899, 95% CI: 0.888,0.911). The linear regression shown in Figure 2B was restricted to ODn values below 6. The Bland-Altman plots in Figure 2C and 2D show that the Maxim assay tends to produce lower OD readings than the Sedia assay on the low end of the dynamic range, and higher readings at the top end. When the calibrators are used to normalize, Maxim ODn values exhibit clear downward bias throughout the dynamic range.

**Figure 2:**
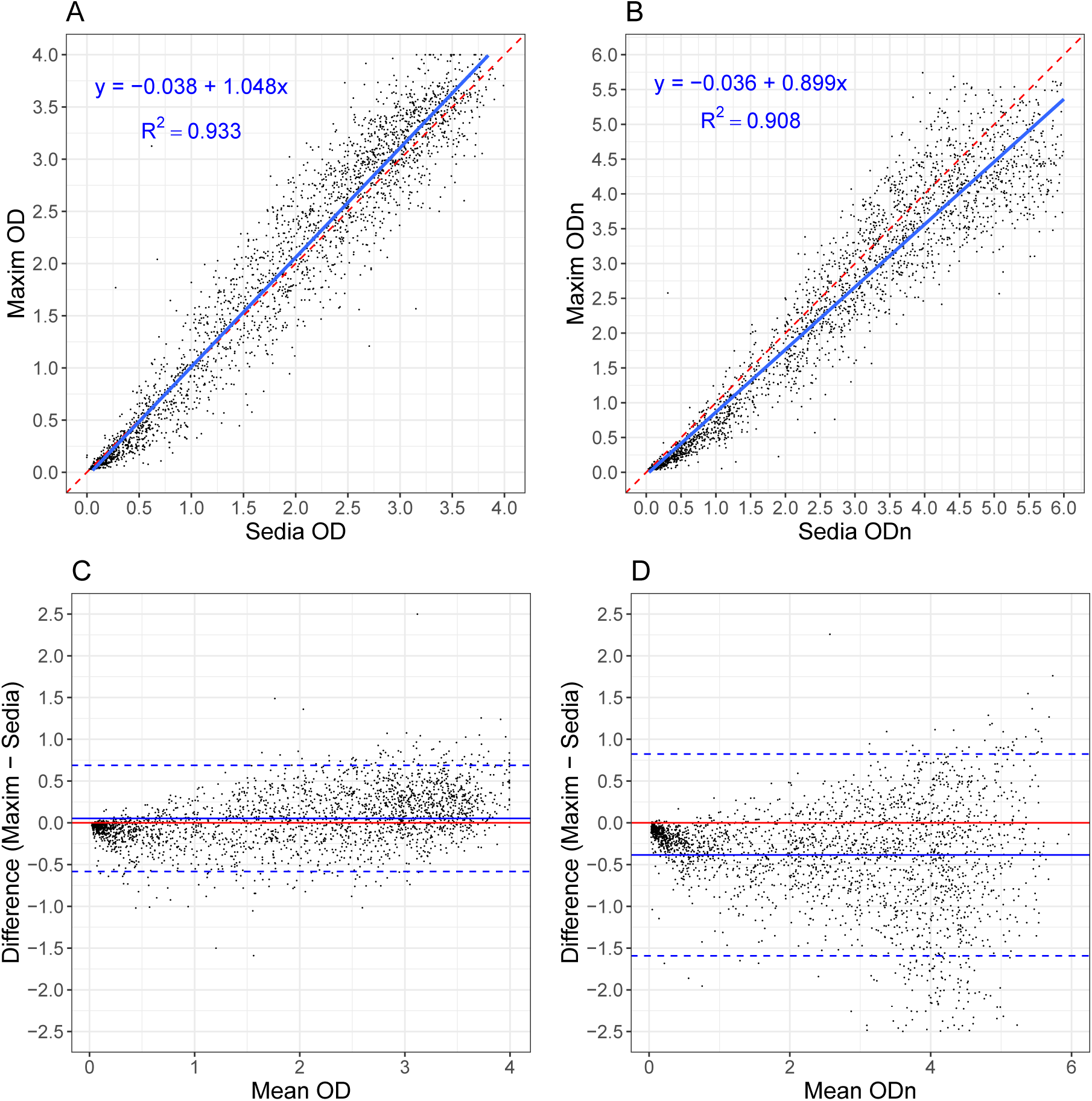
Maxim vs. Sedia OD and ODn measurements. **A.** Maxim vs. Sedia Optical Density (OD); **B.** Maxim vs. Sedia normalized Optical Density (ODn). The blue lines are linear regression fits and the red dashed lines show the diagonal (slope if the two assays produced equivalent results). **C.** Bland-Altman plot for Optical Density (OD); **D.** Bland-Altman plot for normalized Optical Density (ODn). The red lines represent zero bias, the blue solid lines the mean differences and the blue dashed lines the 95% lower and upper limits.

MDRI was estimated using treatment-naïve, non-elite controller subjects, with EDDI intervals ≤120 days. Using ODn ≤1.5, the MDRI for Maxim LAg, without using a supplemental viral load threshold, was 248 days (95% CI: 224,274), while the MDRI for Sedia LAg was 215 days (95% CI: 192,241). This resulted in a statistically significant estimated MDRI difference of 32.7 days (95% CI: 22.9,42.8). Applying a supplemental viral load threshold of >1,000c/mL resulted in MDRI estimates of 201 days (95% CI: 180,224) and 171 days (95% CI: 152,191) for Maxim and Sedia, respectively, with an MDRI difference estimated at 30.9 days (95% CI: 21.7,40.7).

Table 2 shows MDRI estimates for all subtypes, and by subtype (B, C, D and A1), for a range of ODn thresholds in combination with a viral load threshold (>1,000c/mL). We did not observe statistically significant differences between subtype-specific MDRI estimates and the estimates for all other subtypes (using a two-sample Z-test) for either assay at any ODn threshold. MDRI estimates for a wider range of ODn and viral load thresholds are reported in the Appendix (Table A.1).

**Table 2:**
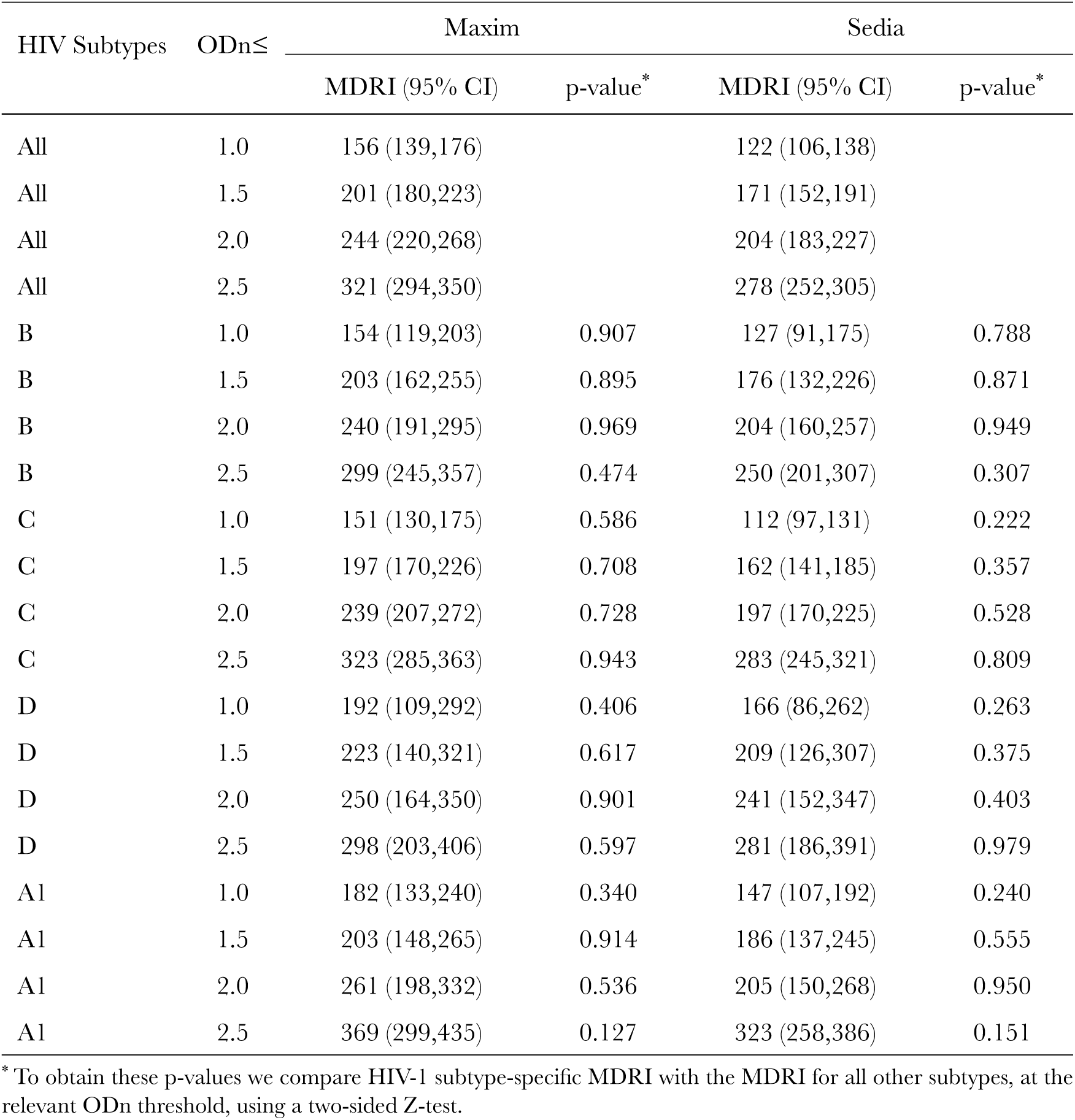
MDRI estimates for Maxim and Sedia LAg assays by HIV-1 subtype and ODn threshold, using supplemental viral load threshold of >1,000c/mL.

The MDRI-against-threshold and (naïvely estimated) FRR-against-threshold trajectories for the two assays were similar as shown in Figures A.1 and A.2 in the Appendix, although the values at any given threshold differed. (The MDRI calibration curve has a characteristic kink at an ODn threshold of 2.0, the threshold that triggers confirmatory testing. This shape is expected, since there are more specimens with higher intrinsic reactivity, which through random fluctuation produce initial ODn values under the threshold and upon confirmatory testing is assigned a final value above—thus depressing the MDRI up to the retesting threshold.)

While naïvely-estimated FRRs at a given threshold were not identical between the Maxim and Sedia assays, the differences were not statistically significant. The FRRs in ART-naïve subjects were 3.26% and 2.17% for Maxim and Sedia, respectively, at ODn ≤1.5, without using supplemental viral load. In combination with a viral load threshold of >1,000c/mL the FRRs were 1.69% and 1.12%, respectively. These estimates are shown in Table A.1 in the Appendix. Among treated subjects FRRs were extremely high when the RITA did not include a viral load threshold. In early-treated subjects (time from infection to treatment initiation ≤6 months), the FRRs for Maxim and Sedia were 98% and 96%, respectively, and in later-treated subjects (time from infection to treatment initiation >6 months), FRRs were 38% vs. 33%, respectively. Using a supplemental viral load threshold reduced these FRRs to 0, given that all treated subjects in the CEPHIA panel were virally suppressed.

### 3.3 Performance in surveillance

The performance of the two assays in the three hypothetical surveillance scenarios defined earlier are summarised in Figures 3 and 4.

**Figure 3:**
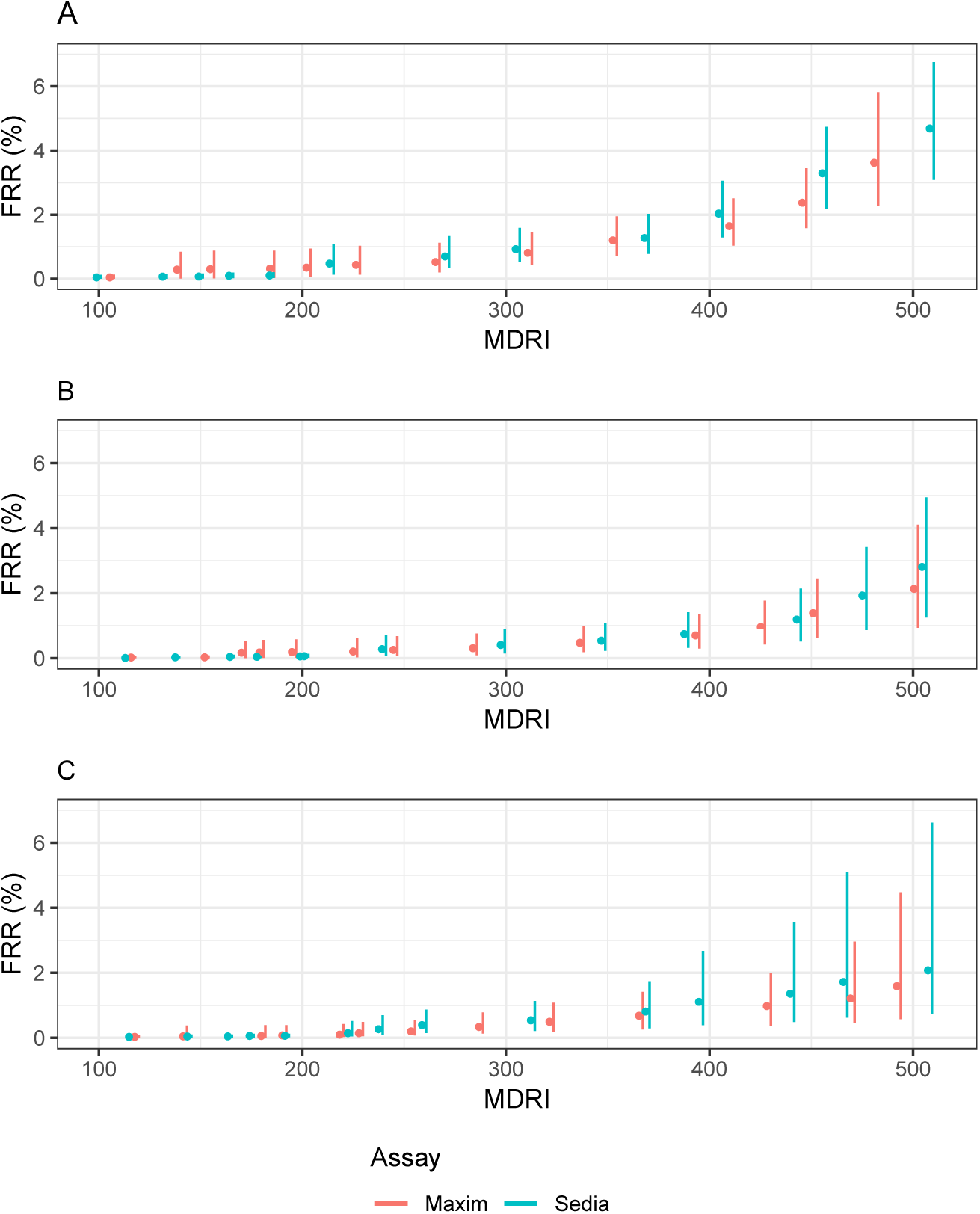
Context-specific false-recent rate (FRR) against MDRI in three demonstrative surveillance scenarios. A supplementary viral load threshold of >1,000c/mL is used throughout. We assume ARV exposure testing classifies all treated individuals as long-term. This assumption is relaxed in Figure A.5. **A.** Scenario similar to South African epidemic. **B.** Scenario similar to Kenyan epidemic. **C.** Scenario similar to North American key population epidemic.

**Figure 4:**
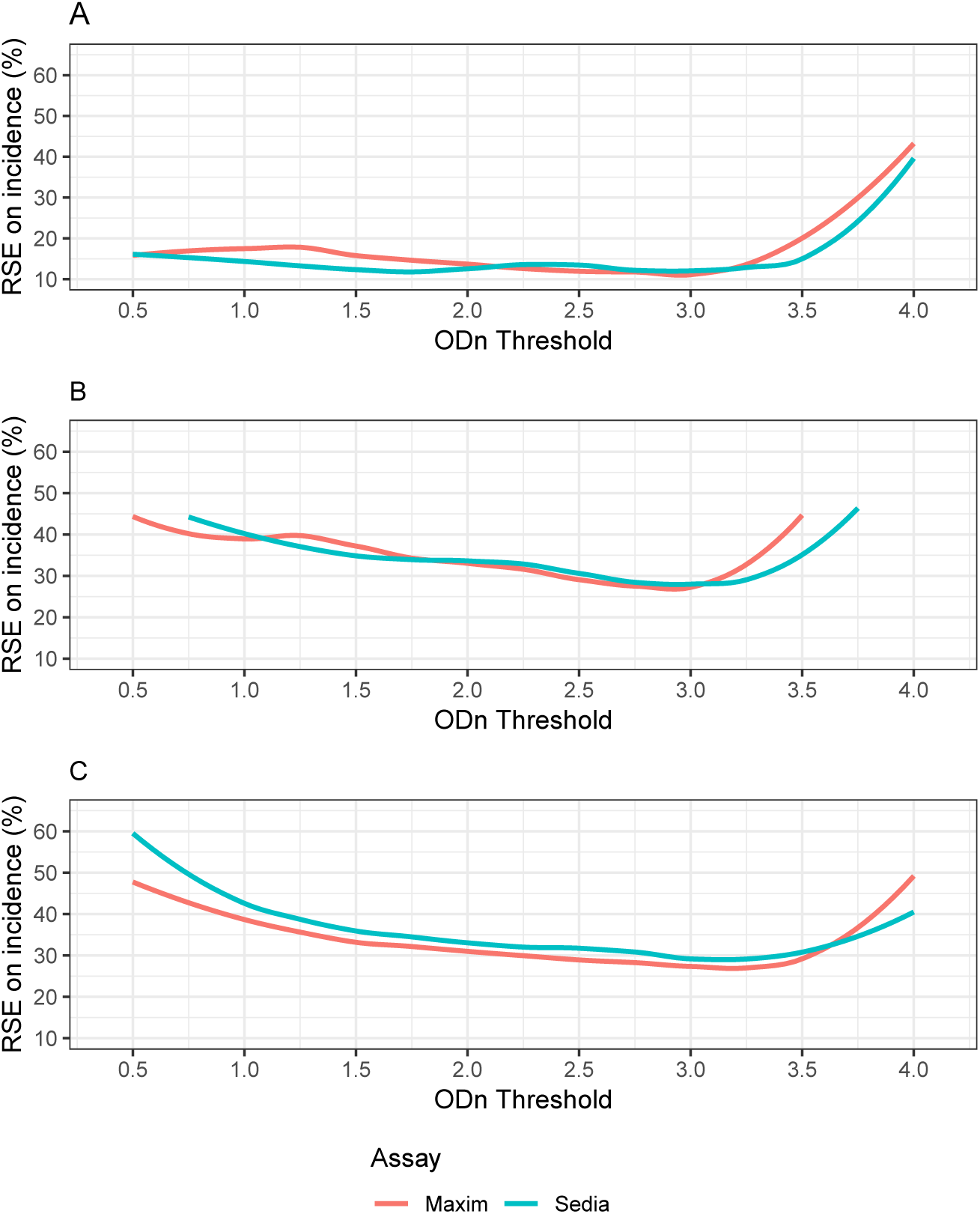
Relative standard error (RSE) of incidence estimate against ODn threshold in three demonstrative surveillance scenarios. A supplementary viral load threshold of >1,000c/mL is used throughout. We assume ARV exposure testing classifies all treated individuals as long-term. This assumption is relaxed in Figure A.6. **A.** Scenario similar to South African epidemic. **B.** Scenario similar to Kenyan epidemic. **C.** Scenario similar to North American key population epidemic.

Figure 3 shows context-specific FRR against MDRI, for RITAs that included a viral load threshold of >1,000c/mL and where we assume that ART exposure testing reduces false recency in treated subjects to zero. Note that the MDRI values on the x-axis encode different ODn thresholds for the two assays. The FRR rises at slightly lower MDRIs for Sedia-based RITAs than for Maxim-based RITAs, in all three scenarios. To maintain FRRs below 2%, both assays require a choice of ODn threshold that produces maximal MDRIs of about 400 to 450 days. In the Appendix, we show context-adapted FRRs against ODn thresholds (Figure A.3) and we relax the assumption that ART exposure testing performs perfectly, instead assuming that it reduces false recency in treated subjects by an order of magnitude compared to that obtained in treated subjects when no supplementary viral load threshold is used. This is shown in Figures A.4 and A.5.

Figure 4 shows the precision of the incidence estimate attained for a range of ODn thresholds in combination with a viral load threshold of >1,000c/mL. At each ODn threshold, assay-specific context-adapted MDRIs and FRRs were computed for use in the incidence calculation. In the South Africa-like scenario, Figure 4A, the lowest value of RSE on incidence attained with the Maxim-based algorithm was 11.7% at ODn ≤3.0, and with the Sedia-based algorithm was 12.0% at the same ODn threshold. In the Kenya-like scenario, Figure 4B, the minimal RSE for Maxim was 27.2%, achieved at ODn ≤2.75, and for Sedia was 28.2% at ODn ≤3.0. In the North American key population-like scenario, Figure 4C, the lowest RSE for Maxim was 26.9% at ODn ≤3.25 and for Sedia was 28.9% at ODn ≤3.0. These nominally optimal thresholds were slightly different under the alternative assumptions shown in the Appendix (Figure A.4). Context-specific MDRI and FRR estimates, and RSEs on incidence estimates, are reported in Table A.2 in the Appendix.

## 4 Discussion

The Maxim and Sedia LAg assays produce meaningfully different ODn results on the same specimens, largely as a result of higher calibrator readings obtained from the Maxim-supplied kit calibrators, and consequently, at any given ODn threshold, RITAs based on the two assays have different MDRIs and FRRs. It is inappropriate to utilize published MDRI and FRR estimates for one assay in survey planning and incidence estimation where the other assay is being used, or to switch from one assay to the other within a study.

It is possible to derive an approximate conversion factor (of 1.172) between ODn values of the two assays from the slopes of the regression curves shown in Figure 2A and 2B. It has further been suggested that a threshold of 1.5 on Maxim is equivalent to a threshold of 2.0 on Sedia, based on testing of a set of specimens, with reactivity spanning the dynamic range, with both assays (personal communication, B. Parekh). Our analysis does indeed show that these thresholds yield very similar MDRIs when used alone (248 days vs. 254 days), but the FRRs are also different. Applying a conversion factor to the Sedia results of repeat-tested specimens does not perfectly predict the Maxim ODn values obtained, and a preferable approach is therefore to use appropriately-estimated MDRIs and FRRs for any given RITA based on either assay.

Our reproducibility analyses show little benefit to the normalization procedure, with both the Maxim and Sedia assays showing greater variability in ODn values than in the raw optical densities on blinded replicate specimens subjected to repeat testing. Further, the correlation between Maxim and Sedia optical densities was greater than between ODn measurements on the same specimens. However, at the time of each of these evaluations, kits and reagents were sourced at the same time, kits were from a small number of lots, and operators were highly experienced with assays. The purpose of the calibrators and normalization procedure is to reduce lot-to-lot variability and ensure stability of results over time and between manufacturers and laboratories. This goal requires that calibrators be highly consistent over time and between manufacturers, which is not currently the case. Fourteen laboratories participate in the NIAID-supported External Quality Assurance Program Oversight Laboratory (EQAPOL) proficiency testing program. This program assessed consistency of results between and within assay manufacturers, kit lots and participating laboratories using panels designed for quality assurance and proficiency testing, and found similar differences in calibrator reactivity and average ODn values between the two assays (Keating et al., forthcoming). External quality assurance is critical for ensuring consistency between laboratories and kit manufacturers.

It should be noted that our evaluation of both assays was restricted to plasma specimens. Both manufacturers also produce kits for use with dried blood spot eluates, and it has been shown that specimen type further impacts performance [16].

We did not observe any statistically significant subtype effects on MDRI, although point estimates differed substantially, especially with specimens from subtype D-infected subjects compared to subtypes B and C (Table 2). With a larger dataset and more precise MDRI estimates, subtype differences may be visible.

Despite the systematic differences in calibrator readings and consequently in the ODn values obtained, performance of the two assays for incidence surveillance was virtually indistinguishable— as long as appropriate assay- and context-specific MDRI and FRR estimates were used. As a result, different ODn thresholds were nominally optimal (i.e. produced the lowest variance on the incidence estimate). In all three hypothetical surveillance scenarios, ODn thresholds between about 1.5 and 3.0 (in combination with viral load), produced the best precision. It is critical, however, that appropriate MDRI and FRR estimates be used for the recency discrimination threshold chosen in order to obtain accurate incidence estimates. It should also be noted that the triplicate ‘confirmatory’ testing protocol mandates confirmatory testing when an initial ODn result is below 2.0, which may be problematic for RITAs that use ODn thresholds above the ‘standard’ threshold of 1.5. It would also be a different subset of specimens reflexed to confirmatory testing on the two assays. Consideration should be given to a modified testing protocol in which confirmatory testing is performed on a larger subset of (or even all) specimens.

A limitation of this study is that we did not have access to specimens from virally unsuppressed treated subjects, and we are therefore unable to rigorously estimate FRR in this group, which may be substantial in many surveillance settings [17]. We urge survey planners and analysts to conduct sensitivity analyses with respect to FRR when utilising either assay in cross-sectional incidence estimation.

Differences in ODn measurements between the Maxim and Sedia LAg assays on the same specimens largely resulted from differences in the reactivity of calibrators supplied by the manufacturers. This resulted in systematically lower ODn measurements on the Maxim assay than on the Sedia assay, and consequently longer MDRIs and larger FRRs at any given ODn recency discrimination threshold. While performance for surveillance purposes was extremely similar, different thresholds were optimal and different values of MDRI and FRR were appropriate for use in survey planning and incidence estimation. The two assays cannot be treated as interchangeable, should not be mixed within one study, and care should be taken when interpreting and comparing results. We summarize our recommendations based on this comparative evaluation in Table 3.

**Table 3:** Summary recommendations for use of the Maxim and Sedia LAg assays.

|  | **Issue** | **Recommendation** |
| --- | --- | --- |
| **Laboratory methods:** | Assay procedures are similar but not identical. | Testing laboratories should ensure full compliance with manufacturer’s instructions for use, especially if both manufacturers’ assays are used in one laboratory. |
| **Quality assurance:** | Lot-to-lot variation and differences in laboratory staff proficiency may further reduce reproducibility of results. | Participation in an external quality assurance programme like EQAPOL [18] is recommended. |
| **Software:** | Although data capture and analysis software are similar, interpretive criteria for specific components differ. | The data analysis software is specific to each assay and laboratories should use the software supplied by the manufacturer. |
| **Conversion:** | Although it is possible to compute an approximate conversion factor, this does not perfectly predict equivalent ODn values. | Rather than converting results, appropriately-derived MDRI and FRR estimates should be utilized for each assay. The same ODn thresholds may not be optimal. |
| **Descriptive title:** | The names ‘HIV-1 Limiting Antigen Avidity EIA’ or ‘LAg assay’ do not distinguish between the two assays. | Users should clearly identify the manufacturer of the kits used, as well as specimen type, in all publications and reports. |
| **Assay performance:** | Despite differences in calibrator reactivity, and consequently in ODn values obtained on the same specimens, performance of the two assays for surveillance purposes was virtually indistinguishable. | Both manufacturers’ assays are suitable for use, but they should not be mixed within studies, appropriate performance characteristic estimates must be used and care should be taken when comparing results. |

## Author contributions

J.B.S. and E.G. analysed the data; J.B.S., G.M. and E.G. wrote the article; G.M. managed the laboratory testing; J.H. conducted laboratory testing; S.N.F., D.H., S.M.K., K.M. and E.G. managed the specimen repository and clinical data; N.P. provided scientific support and managed the primary grant; G.M., A.W., C.D.P., and M.P.B. were principal investigators for the study and provided input on analysis and writing; All authors reviewed and provided input on the article.

## Acknowledgments

The authors gratefully acknowledge the Centre for High Performance Computing, which provided computational resources for this study.

The Consortium for the Evaluation and Performance of HIV Incidence Assays (CEPHIA) comprises: Oliver Laeyendecker, Thomas Quinn, David Burns (National Institutes of Health); Alex Welte, Joseph Sempa, previously: David Matten, Hilmarié Brand, Trust Chibawara (South African Centre for Epidemiological Modelling and Analysis); Reshma Kassanjee (University of Cape Town); Gary Murphy, Jake Hall, previously: Elaine Mckinney (Public Health England); Michael P. Busch, Eduard Grebe, Shelley Facente, Sheila Keating, Dylan Hampton, previously: Mila Lebedeva (Vitalant Research Institute, formerly Blood Systems Research Institute); Christopher D. Pilcher, Kara Marson (University of California, San Francisco); Susan Little (University of California, San Diego); Anita Sands (World Health Organization); Tim Hallett (Imperial College London); Sherry Michele Owen, Bharat Parekh, Connie Sexton (Centers for Disease Control and Prevention); Matthew Price, Anatoli Kamali (International AIDS Vaccine Initiative); Lisa Loeb (The Options Study—University of California, San Francisco); Jeffrey Martin, Steven G Deeks, Rebecca Hoh (The SCOPE Study— University of California, San Francisco); Zelinda Bartolomei, Natalia Cerqueira (The AMPLIAR Cohort—University of São Paulo); Breno Santos, Kellin Zabtoski, Rita de Cassia Alves Lira (The AMPLIAR Cohort—Grupo Hospital Conceição); Rosa Dea Sperhacke, Leonardo R Motta, Machline Paganella (The AMPLIAR Cohort—Universidade Caxias Do Sul); Esper Kallas, Helena Tomiyama, Claudia Tomiyama, Priscilla Costa, Maria A Nunes, Gisele Reis, Mariana M Sauer, Natalia Cerqueira, Zelinda Nakagawa, Lilian Ferrari, Ana P Amaral, Karine Milani (The São Paulo Cohort—University of São Paulo, Brazil); Salim S Abdool Karim, Quarraisha Abdool Karim, Thumbi Ndungu, Nelisile Majola, Natasha Samsunder (CAPRISA, University of Kwazulu-Natal); Denise Naniche (The GAMA Study—Barcelona Centre for International Health Research); Inácio Mandomando, Eusebio V Macete (The GAMA Study—Fundacao Manhica); Jorge Sanchez, Javier Lama (SABES Cohort—Asociación Civil Impacta Salud y Educación (IMPACTA)); Ann Duerr (The Fred Hutchinson Cancer Research Center); Maria R Capobianchi (National Institute for Infectious Diseases “L. Spallanzani”, Rome); Barbara Suligoi (Istituto Superiore di Sanità, Rome); Susan Stramer (American Red Cross); Phillip Williamson (Creative Testing Solutions / Vitalant Research Institute); Marion Vermeulen (South African National Blood Service); and Ester Sabino (Hemocentro do São Paolo). General inquiries may be directed to

## A Appendix

**Table A.1:** MDRI and FRR* estimates in ART-naïve subjects for a range of ODn and viral load thresholds.

| ODn<br>≤ | VL<br>> | Maxim |  | Sedia |  |
| --- | --- | --- | --- | --- | --- |
|  |  | MDRI (95% CI) | FRR (95% CI) | MDRI (95% CI) | FRR (95% CI) |
| 0.50 | None | 129 (113,146) | 1.6% (0.3,4.7) | 91 (79,105) | 0.5% (0.0,3.0) |
| 0.75 | None | 164 (144,184) | 1.6% (0.3,4.7) | 127 (111,145) | 1.6% (0.3,4.7) |
| 1.00 | None | 197 (176,220) | 2.2% (0.6,5.5) | 158 (138,179) | 1.6% (0.3,4.7) |
| 1.25 | None | 224 (200,248) | 2.7% (0.9,6.2) | 194 (172,217) | 2.2% (0.6,5.5) |
| 1.50 | None | 248 (224,274) | 3.3% (1.2,7.0) | 215 (192,240) | 2.2% (0.6,5.5) |
| 1.75 | None | 277 (251,304) | 3.8% (1.5,7.7) | 237 (213,263) | 4.4% (1.9,8.4) |
| 2.00 | None | 297 (270,324) | 6.5% (3.4,11.1) | 254 (228,280) | 4.4% (1.9,8.4) |
| 2.25 | None | 338 (311,365) | 7.6% (4.2,12.4) | 292 (265,319) | 6.0% (3.0,10.4) |
| 2.50 | None | 382 (354,410) | 11.4% (7.2,16.9) | 333 (305,362) | 9.8% (5.9,15.0) |
| 2.75 | None | 423 (394,453) | 16.9% (11.7,23.1) | 373 (344,401) | 13.0% (8.5,18.8) |
| 3.00 | None | 471 (442,500) | 20.1% (14.6,26.6) | 427 (398,457) | 16.3% (11.3,22.5) |
| 0.50 | 0 | 129 (113,147) | 1.7% (0.4,4.9) | 91 (79,105) | 0.6% (0.0,3.1) |
| 0.50 | 75 | 118 (104,132) | 1.1% (0.1,4.0) | 83 (73,94) | 0.6% (0.0,3.1) |
| 0.50 | 400 | 108 (96,122) | 0.6% (0.0,3.1) | 78 (68,88) | 0.0% (0.0,2.1) |
| 0.50 | 1,000 | 102 (90,114) | 0.6% (0.0,3.1) | 73 (64,82) | 0.0% (0.0,2.1) |
| 0.50 | 5,000 | 86 (75,97) | 0.6% (0.0,3.1) | 63 (54,72) | 0.0% (0.0,2.1) |
| 0.75 | 0 | 165 (145,186) | 1.7% (0.4,4.9) | 127 (110,146) | 1.7% (0.4,4.9) |
| 0.75 | 75 | 151 (133,169) | 1.1% (0.1,4.0) | 116 (102,132) | 0.6% (0.0,3.1) |
| 0.75 | 400 | 138 (121,155) | 0.6% (0.0,3.1) | 106 (93,120) | 0.6% (0.0,3.1) |
| 0.75 | 1,000 | 127 (112,144) | 0.6% (0.0,3.1) | 99 (86,112) | 0.6% (0.0,3.1) |
| 0.75 | 5,000 | 106 (92,122) | 0.6% (0.0,3.1) | 83 (71,96) | 0.6% (0.0,3.1) |
| 1.00 | 0 | 199 (177,223) | 2.3% (0.6,5.7) | 159 (139,181) | 1.7% (0.4,4.9) |
| 1.00 | 75 | 183 (164,205) | 1.1% (0.1,4.0) | 145 (127,164) | 1.1% (0.1,4.0) |
| 1.00 | 400 | 168 (149,188) | 1.1% (0.1,4.0) | 130 (114,148) | 0.6% (0.0,3.1) |
| 1.00 | 1,000 | 156 (139,176) | 1.1% (0.1,4.0) | 122 (106,138) | 0.6% (0.0,3.1) |
| 1.00 | 5,000 | 126 (109,144) | 1.1% (0.1,4.0) | 101 (86,117) | 0.6% (0.0,3.1) |
| 1.25 | 0 | 226 (202,251) | 2.8% (0.9,6.4) | 197 (174,221) | 2.3% (0.6,5.7) |
| 1.25 | 75 | 210 (188,233) | 1.7% (0.4,4.9) | 180 (161,201) | 1.7% (0.4,4.9) |
| 1.25 | 400 | 191 (171,212) | 1.7% (0.4,4.9) | 164 (146,184) | 1.1% (0.1,4.0) |
| 1.25 | 1,000 | 179 (160,199) | 1.7% (0.4,4.9) | 153 (135,172) | 1.1% (0.1,4.0) |
| 1.25 | 5,000 | 141 (123,160) | 1.7% (0.4,4.9) | 122 (106,140) | 1.1% (0.1,4.0) |
| 1.50 | 0 | 250 (225,276) | 3.4% (1.3,7.2) | 218 (194,244) | 2.3% (0.6,5.7) |
| 1.50 | 75 | 233 (210,256) | 1.7% (0.4,4.9) | 202 (180,224) | 1.7% (0.4,4.9) |
| 1.50 | 400 | 214 (193,237) | 1.7% (0.4,4.9) | 183 (163,204) | 1.1% (0.1,4.0) |
| 1.50 | 1,000 | 201 (180,223) | 1.7% (0.4,4.9) | 171 (152,191) | 1.1% (0.1,4.0) |
| 1.50 | 5,000 | 158 (138,179) | 1.7% (0.4,4.9) | 135 (118,154) | 1.1% (0.1,4.0) |
| 1.75 | 0 | 279 (252,306) | 3.9% (1.6,7.9) | 239 (213,266) | 4.5% (2.0,8.7) |
| 1.75 | 75 | 261 (236,286) | 1.7% (0.4,4.9) | 222 (199,245) | 2.3% (0.6,5.7) |
| 1.75 | 400 | 242 (219,266) | 1.7% (0.4,4.9) | 203 (182,225) | 1.7% (0.4,4.9) |
| 1.75 | 1,000 | 227 (206,251) | 1.7% (0.4,4.9) | 190 (170,211) | 1.7% (0.4,4.9) |
| 1.75 | 5,000 | 175 (155,198) | 1.7% (0.4,4.9) | 148 (129,168) | 1.7% (0.4,4.9) |
| 2.00 | 0 | 299 (272,326) | 6.7% (3.5,11.5) | 254 (228,282) | 4.5% (2.0,8.7) |
| 2.00 | 75 | 281 (255,307) | 5.1% (2.3,9.4) | 236 (212,261) | 2.8% (0.9,6.4) |
| 2.00 | 400 | 261 (237,286) | 3.9% (1.6,7.9) | 217 (196,241) | 1.7% (0.4,4.9) |
| 2.00 | 1,000 | 244 (220,268) | 3.9% (1.6,7.9) | 204 (183,227) | 1.7% (0.4,4.9) |
| 2.00 | 5,000 | 188 (166,210) | 3.9% (1.6,7.9) | 160 (141,180) | 1.7% (0.4,4.9) |
| 2.25 | 0 | 339 (312,367) | 7.9% (4.4,12.8) | 292 (265,320) | 6.2% (3.1,10.8) |
| 2.25 | 75 | 321 (295,348) | 6.2% (3.1,10.8) | 274 (249,300) | 4.5% (2.0,8.7) |
| 2.25 | 400 | 300 (276,326) | 4.5% (2.0,8.7) | 253 (230,278) | 3.4% (1.3,7.2) |
| 2.25 | 1,000 | 280 (255,306) | 4.5% (2.0,8.7) | 237 (214,261) | 3.4% (1.3,7.2) |
| 2.25 | 5,000 | 214 (190,239) | 3.9% (1.6,7.9) | 186 (164,209) | 2.8% (0.9,6.4) |
| 2.50 | 0 | 383 (354,411) | 11.2% (7.0,16.8) | 335 (306,364) | 10.1% (6.1,15.5) |
| 2.50 | 75 | 364 (336,392) | 9.6% (5.7,14.9) | 316 (289,344) | 8.4% (4.8,13.5) |
| 2.50 | 400 | 343 (316,371) | 7.9% (4.4,12.8) | 296 (269,322) | 6.7% (3.5,11.5) |
| 2.50 | 1,000 | 321 (294,350) | 7.9% (4.4,12.8) | 278 (252,305) | 6.7% (3.5,11.5) |
| 2.50 | 5,000 | 252 (225,280) | 6.2% (3.1,10.8) | 220 (194,246) | 5.1% (2.3,9.4) |
| 2.75 | 0 | 426 (397,455) | 17.4% (12.2,23.8) | 375 (346,405) | 13.5% (8.8,19.4) |
| 2.75 | 75 | 406 (377,434) | 15.7% (10.7,21.9) | 357 (328,386) | 11.8% (7.5,17.5) |
| 2.75 | 400 | 384 (355,413) | 13.5% (8.8,19.4) | 336 (308,364) | 10.1% (6.1,15.5) |
| 2.75 | 1,000 | 363 (334,393) | 13.5% (8.8,19.4) | 315 (287,343) | 9.6% (5.7,14.9) |
| 2.75 | 5,000 | 285 (256,315) | 10.7% (6.6,16.2) | 250 (223,279) | 6.2% (3.1,10.8) |
| 3.00 | 0 | 475 (445,505) | 20.8% (15.1,27.5) | 431 (401,461) | 16.9% (11.7,23.2) |
| 3.00 | 75 | 455 (426,483) | 19.1% (13.6,25.7) | 412 (382,441) | 15.2% (10.2,21.3) |
| 3.00 | 400 | 430 (401,459) | 16.9% (11.7,23.2) | 391 (362,420) | 13.5% (8.8,19.4) |
| 3.00 | 1,000 | 408 (379,437) | 16.9% (11.7,23.2) | 369 (340,396) | 13.5% (8.8,19.4) |
| 3.00 | 5,000 | 326 (297,358) | 12.9% (8.4,18.8) | 296 (266,327) | 9.6% (5.7,14.9) |

\*Naïvely-estimated FRR, i.e. not adapted to epidemiological context.

**Table A.2:** Context-specific MDRI and FRR estimates from the three demonstrative surveillance scenarios under different assumptions about impact of ARV exposure testing on FRR.

| Assay | ODn | Scenario<sup>§</sup> | MDRI<sup>†</sup><br>(95% CI) | Assumption 1<sup>*</sup> |  | Assumption 2<sup>**</sup> |  |
| --- | --- | --- | --- | --- | --- | --- | --- |
|  |  |  |  | FRR (95% CI) | RSE | FRR (95% CI) | RSE |
| Maxim | 0.5 | A | 88 (74,104) | 0.0% (0.0,0.1) | 16.7% | 0.6% (0.1,2.4) | 63.8% |
| Maxim | 1.0 | A | 140 (119,165) | 0.3% (0.0,0.8) | 19.0% | 0.9% (0.1,3.0) | 46.8% |
| Maxim | 1.5 | A | 186 (159,214) | 0.3% (0.0,0.9) | 15.6% | 1.0% (0.2,3.3) | 38.1% |
| Maxim | 2.0 | A | 228 (196,261) | 0.4% (0.1,1.0) | 13.8% | 1.2% (0.3,3.7) | 34.2% |
| Maxim | 2.5 | A | 313 (273,353) | 0.8% (0.4,1.5) | 11.8% | 1.7% (0.7,4.4) | 27.3% |
| Sedia | 0.5 | A | 73 (61,85) | 0.0% (0.0,0.0) | 16.1% | 0.5% (0.0,2.0) | 63.5% |
| Sedia | 1.0 | A | 101 (86,120) | 0.0% (0.0,1.0) | 14.7% | 0.7% (0.1,2.6) | 58.9% |
| Sedia | 1.5 | A | 151 (130,174) | 0.1% (0.0,0.2) | 12.3% | 0.8% (0.1,2.8) | 42.1% |
| Sedia | 2.0 | A | 186 (159,215) | 0.1% (0.0,0.2) | 11.7% | 0.8% (0.1,3.0) | 36.5% |
| Sedia | 2.5 | A | 272 (234,311) | 0.7% (0.3,1.3) | 13.1% | 1.5% (0.6,4.1) | 30.0% |
| Maxim | 0.5 | B | 88 (74,104) | 0.0% (0.0,0.1) | 16.7% | 0.6% (0.1,2.4) | 63.8% |
| Maxim | 1.0 | B | 140 (119,165) | 0.3% (0.0,0.8) | 19.0% | 0.9% (0.1,2.0) | 46.8% |
| Maxim | 1.5 | B | 186 (159,214) | 0.3% (0.0,0.9) | 15.6% | 1.0% (0.2,3.3) | 38.1% |
| Maxim | 2.0 | B | 228 (196,261) | 0.4% (0.1,1.0) | 13.8% | 1.2% (0.3,3.7) | 34.2% |
| Maxim | 2.5 | B | 313 (273,353) | 0.8% (0.4,1.5) | 11.8% | 1.7% (0.7,4.4) | 27.3% |
| Sedia | 0.5 | B | 73 (61,85) | 0.0% (0.0,0.0) | 16.1% | 0.5% (0.0,0.2) | 63.5% |
| Sedia | 1.0 | B | 101 (86,120) | 0.0% (0.0,0.1) | 14.7% | 0.7% (0.1,2.6) | 58.9% |
| Sedia | 1.5 | B | 151 (130,174) | 0.1% (0.0,0.2) | 12.3% | 0.8% (0.1,2.8) | 42.1% |
| Sedia | 2.0 | B | 186 (159,215) | 0.1% (0.0,0.2) | 11.7% | 0.8% (0.1,3.0) | 36.5% |
| Sedia | 2.5 | B | 272 (234,311) | 0.7% (0.3,1.3) | 13.1% | 1.5% (0.6,4.1) | 29.9% |
| Maxim | 0.5 | C | 88 (74,104) | 0.0% (0.0,0.1) | 16.7% | 0.6% (0.1,2.4) | 63.8% |
| Maxim | 1.0 | C | 140 (119,165) | 0.3% (0.0,0.8) | 19.0% | 0.9% (0.1,3.0) | 46.8% |
| Maxim | 1.5 | C | 186 (159,214) | 0.3% (0.0,0.9) | 15.6% | 1.0% (0.2,3.3) | 38.1% |
| Maxim | 2.0 | C | 228 (196,261) | 0.4% (0.1,1.0) | 13.8% | 1.2% (0.3,3.7) | 34.2% |
| Maxim | 2.5 | C | 313 (273,353) | 0.8% (0.4,1.5) | 11.8% | 1.7% (0.7,4.4) | 27.3% |
| Sedia | 0.5 | C | 73 (61,85) | 0.0% (0.0,0.0) | 16.1% | 0.5% (0.0,2.0) | 63.5% |
| Sedia | 1.0 | C | 101 (86,120) | 0.0% (0.0,0.1) | 14.7% | 0.7% (0.1,2.6) | 58.9% |
| Sedia | 1.5 | C | 151 (130,174) | 0.1% (0.0,0.2) | 12.3% | 0.8% (0.1,2.8) | 42.0% |
| Sedia | 2.0 | C | 186 (159,215) | 0.1% (0.0,0.2) | 11.7% | 0.8% (0.1,3.0) | 36.5% |
| Sedia | 2.5 | C | 272 (234,311) | 0.7% (0.3,1.3) | 13.1% | 1.5% (0.6,4.1) | 29.9% |

<sup>§</sup>Scenario **A.** South Africa-like epidemic; **B.** Kenya-like epidemic; **C.** North American key population-like epidemic.
<sup>†</sup>Adjusted for subtype mix and screening assay
<sup>\*</sup>ARV exposure testing classifies all treated subjects as long-term
<sup>\*\*</sup>ARV exposure testing reduces FRR in treated subjects to 10% of FRR without viral load

**Figure A.1:**
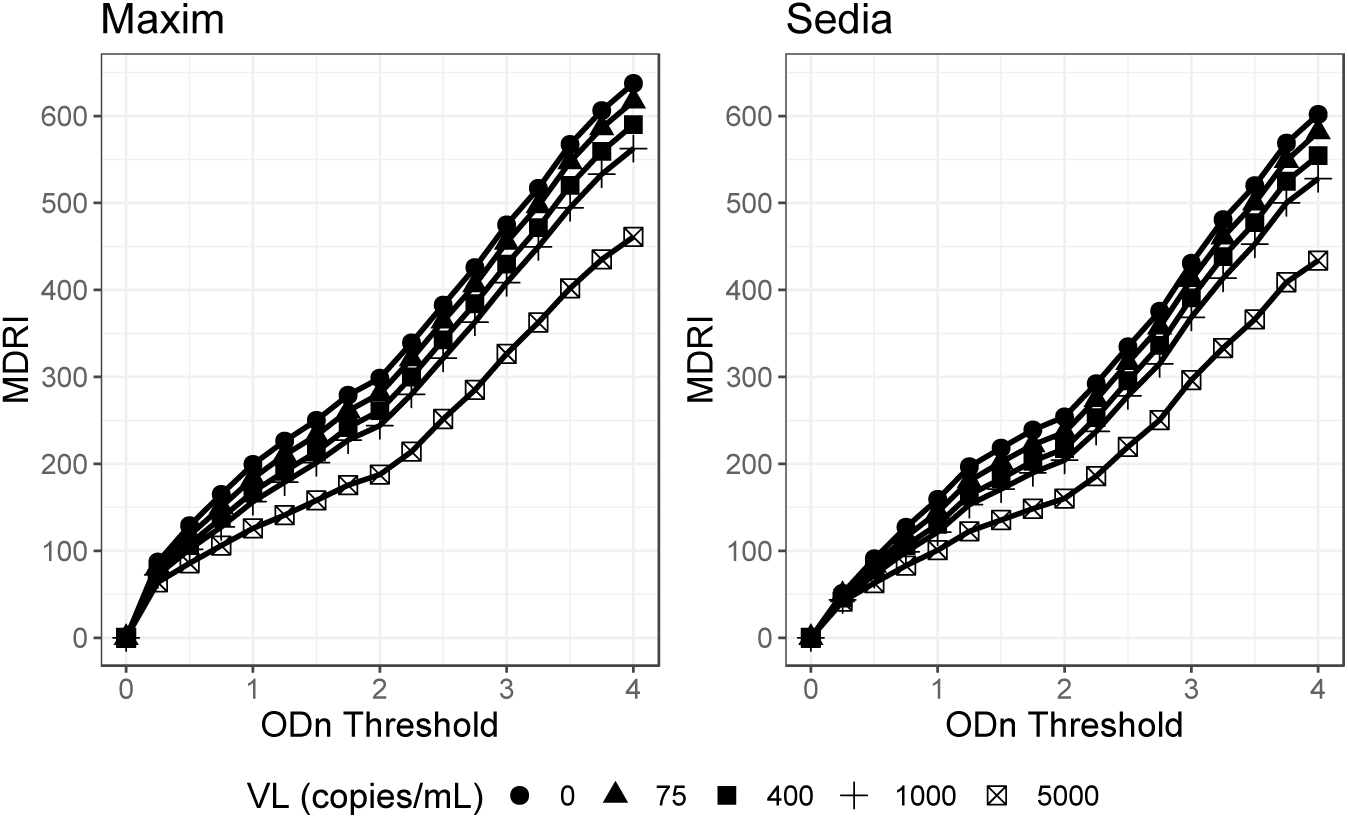
MDRI point estimates against ODn thresholds at different supplemental viral load thresholds. **A.** Maxim; **B.** Sedia

**Figure A.2:**
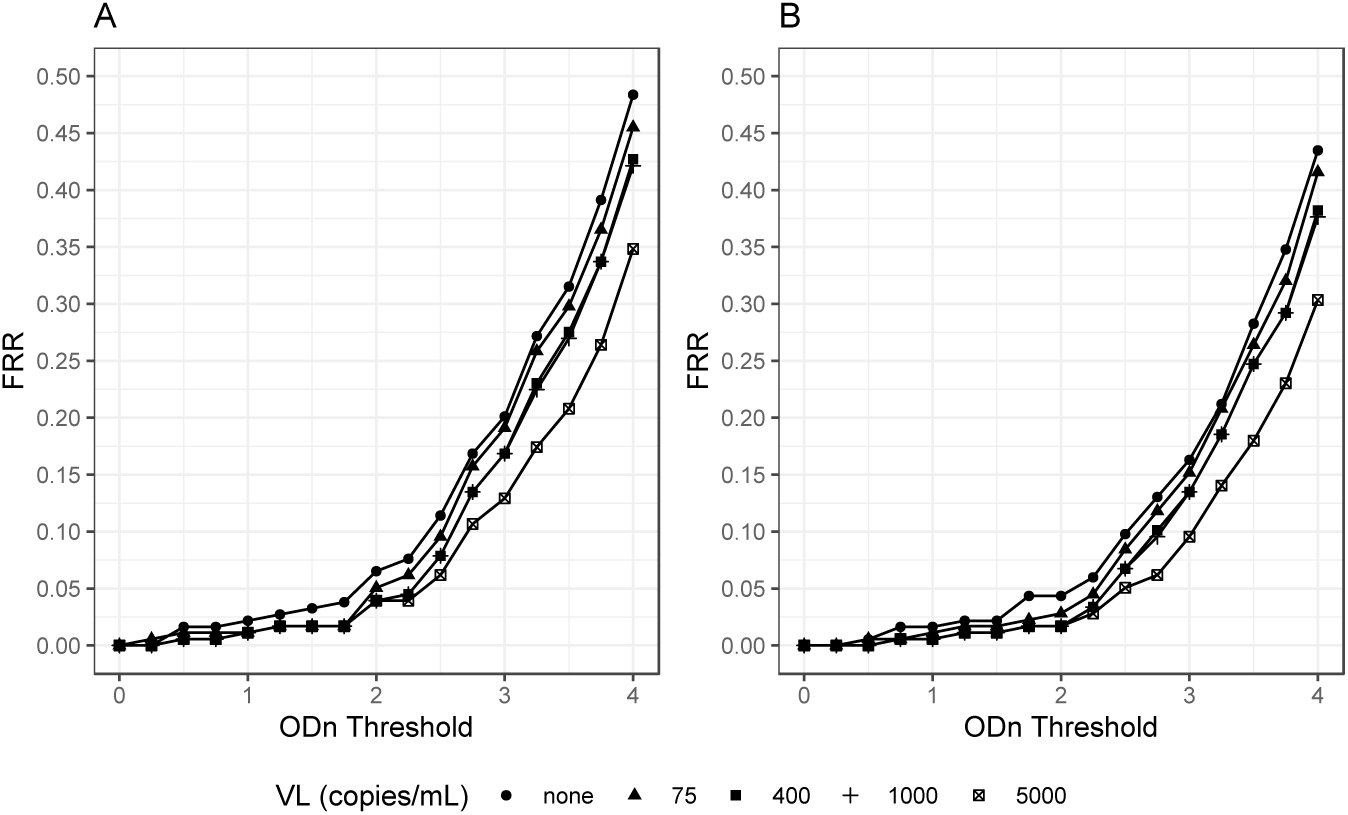
FRR estimates in ART-naïve patients against ODn threshold at different supplemental viral load thresholds. **A.** Maxim; **B.** Sedia

**Figure A.3:**
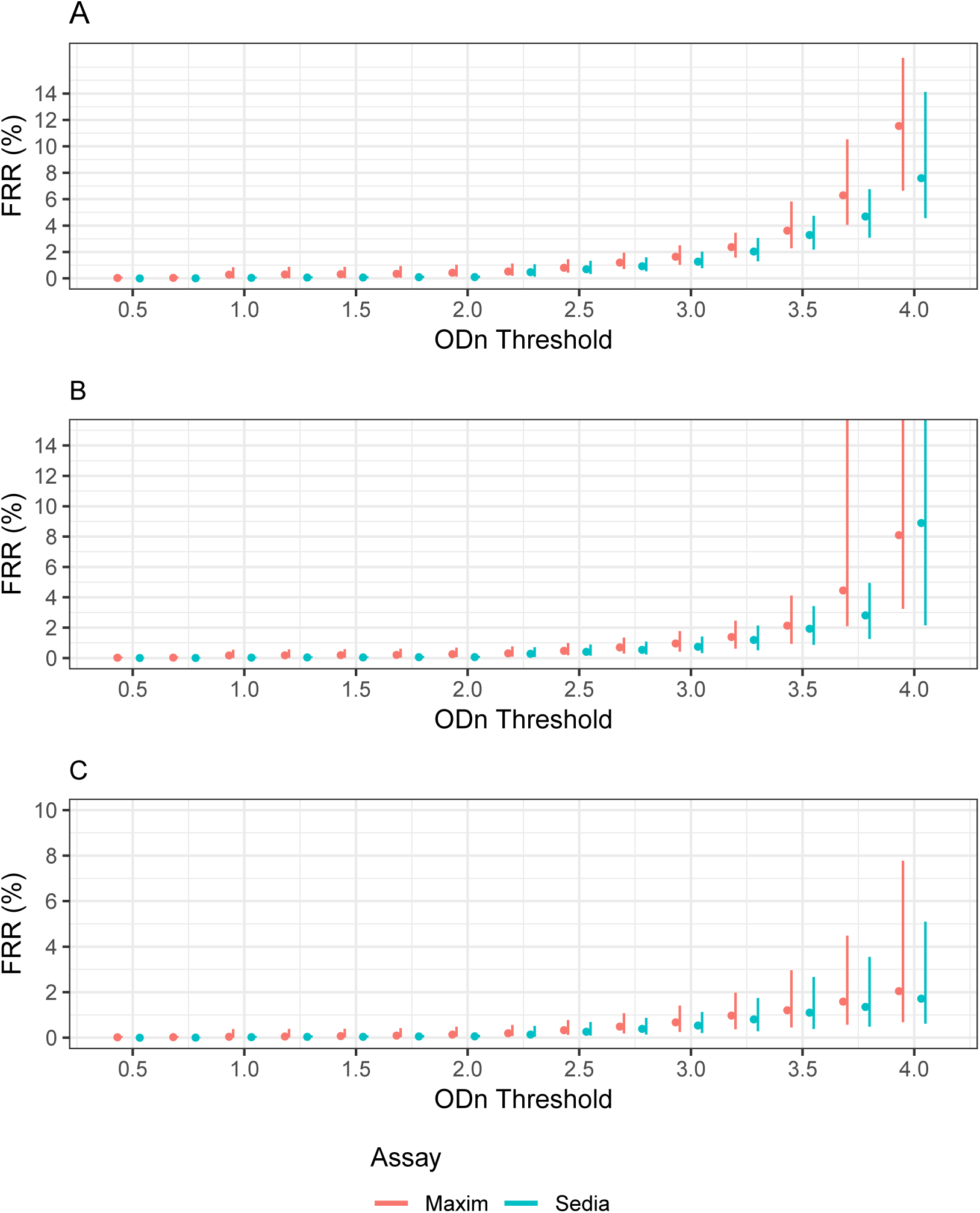
Context-specific FRR vs. ODn threshold in three demonstrative surveillance scenarios (assuming perfect ARV exposure testing) A supplementary viral load threshold of >1,000c/mL is used throughout. We assume ARV exposure testing classifies all treated individuals as long-term. **A.** Scenario similar to South African epidemic. **B.** Scenario similar to Kenyan epidemic. **C.** Scenario similar to North American key population epidemic.

**Figure A.4:**
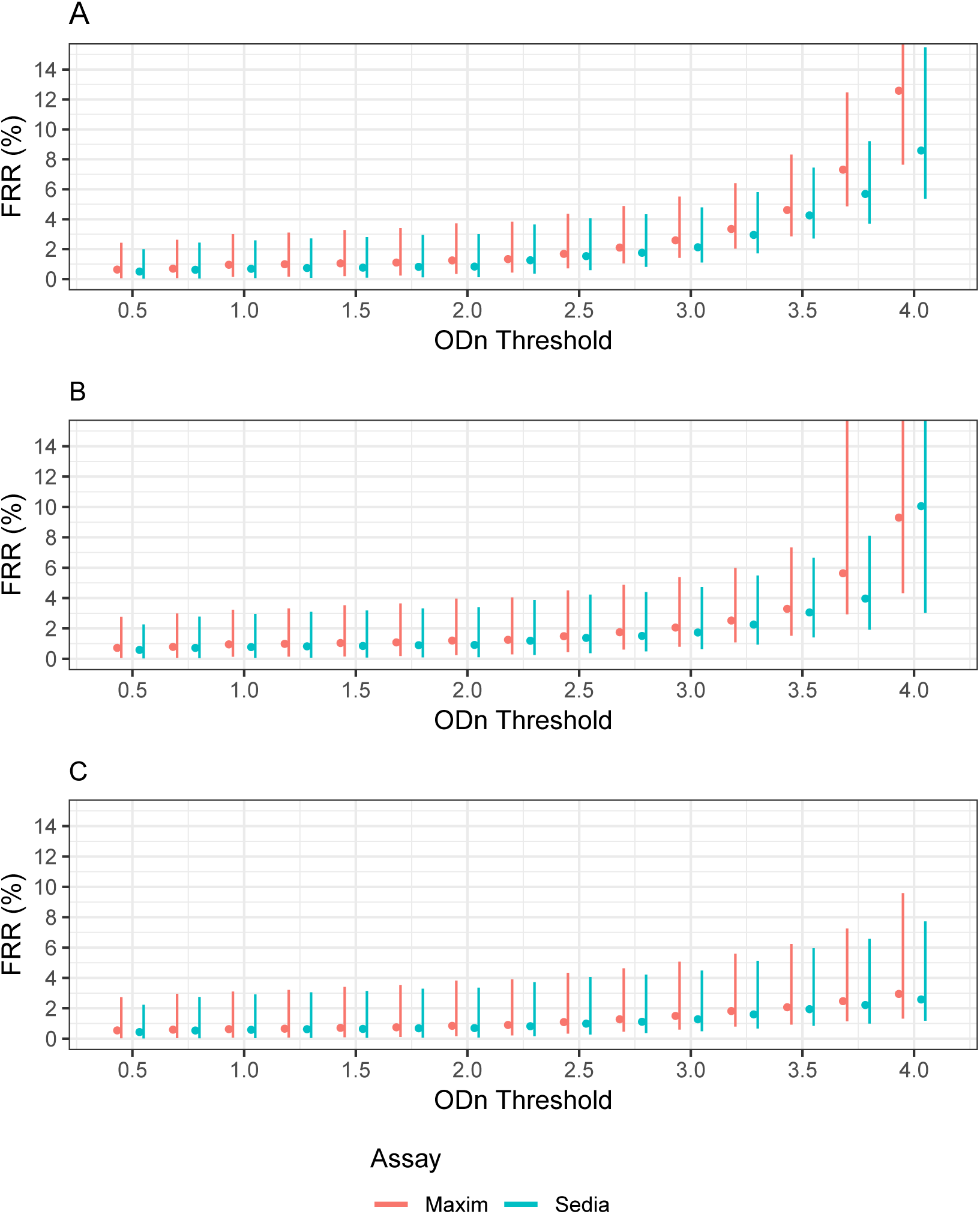
Context-specific FRR vs. ODn threshold in three demonstrative surveillance scenarios (assuming imperfect ARV exposure testing) A supplementary viral load threshold of >1,000c/mL is used throughout. We assume ARV exposure testing reduces false recency in treated individuals to 10% of that attained when no supplemental viral load threshold is utilized. **A.** Scenario similar to South African epidemic. **B.** Scenario similar to Kenyan epidemic. **C.** Scenario similar to North American key population epidemic.

**Figure A.5:**
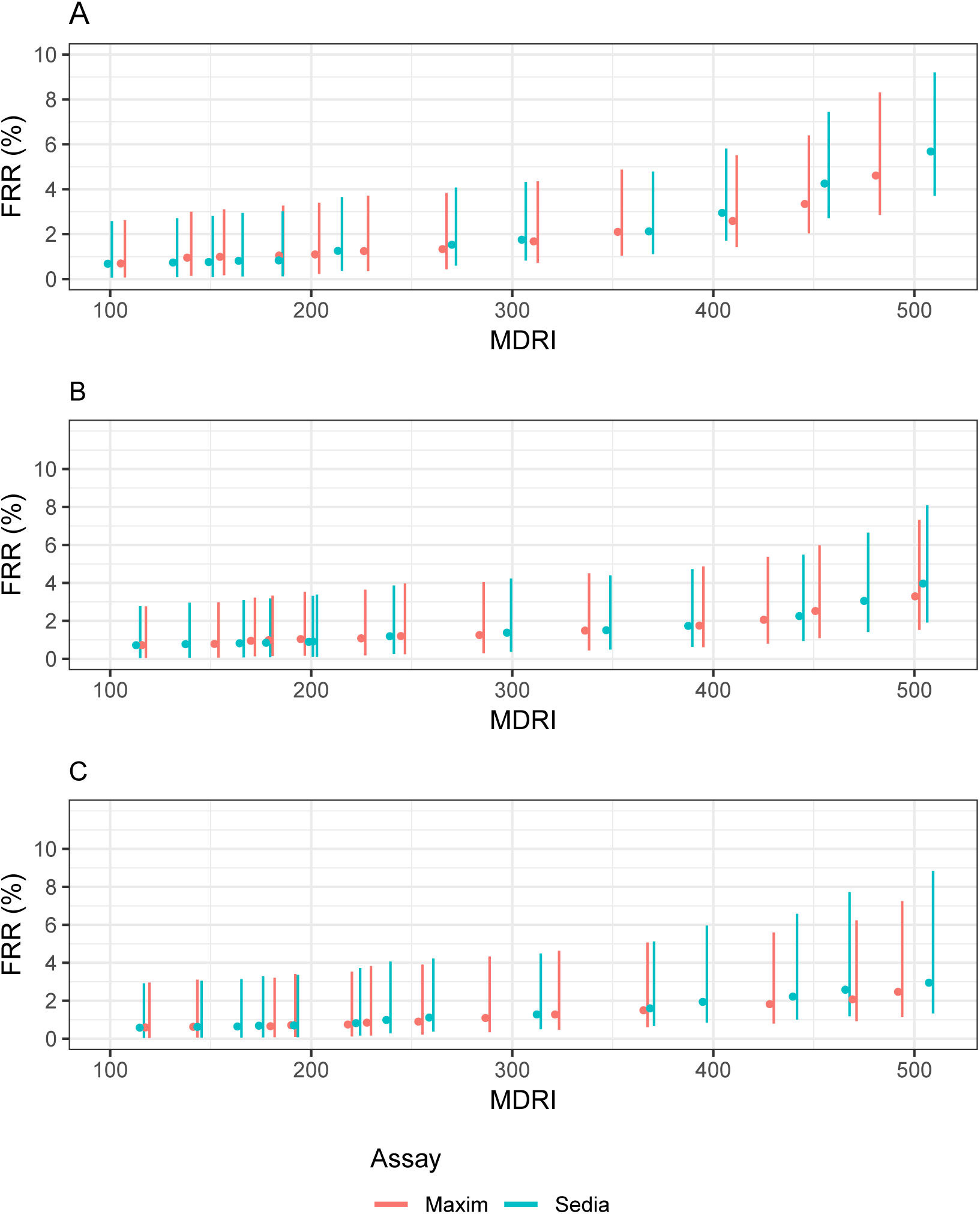
Context-specific false-recent rate (FRR) against MDRI in three demonstrative surveillance scenarios (assuming imperfect ARV testing) A supplementary viral load threshold of >1,000c/mL is used throughout. We assume ARV exposure testing reduces false recency in treated individuals to 10% of that attained when no supplemental viral load threshold is utilized. **A.** Scenario similar to South African epidemic. **B.** Scenario similar to Kenyan epidemic. **C.** Scenario similar to North American key population epidemic.

**Figure A.6:**
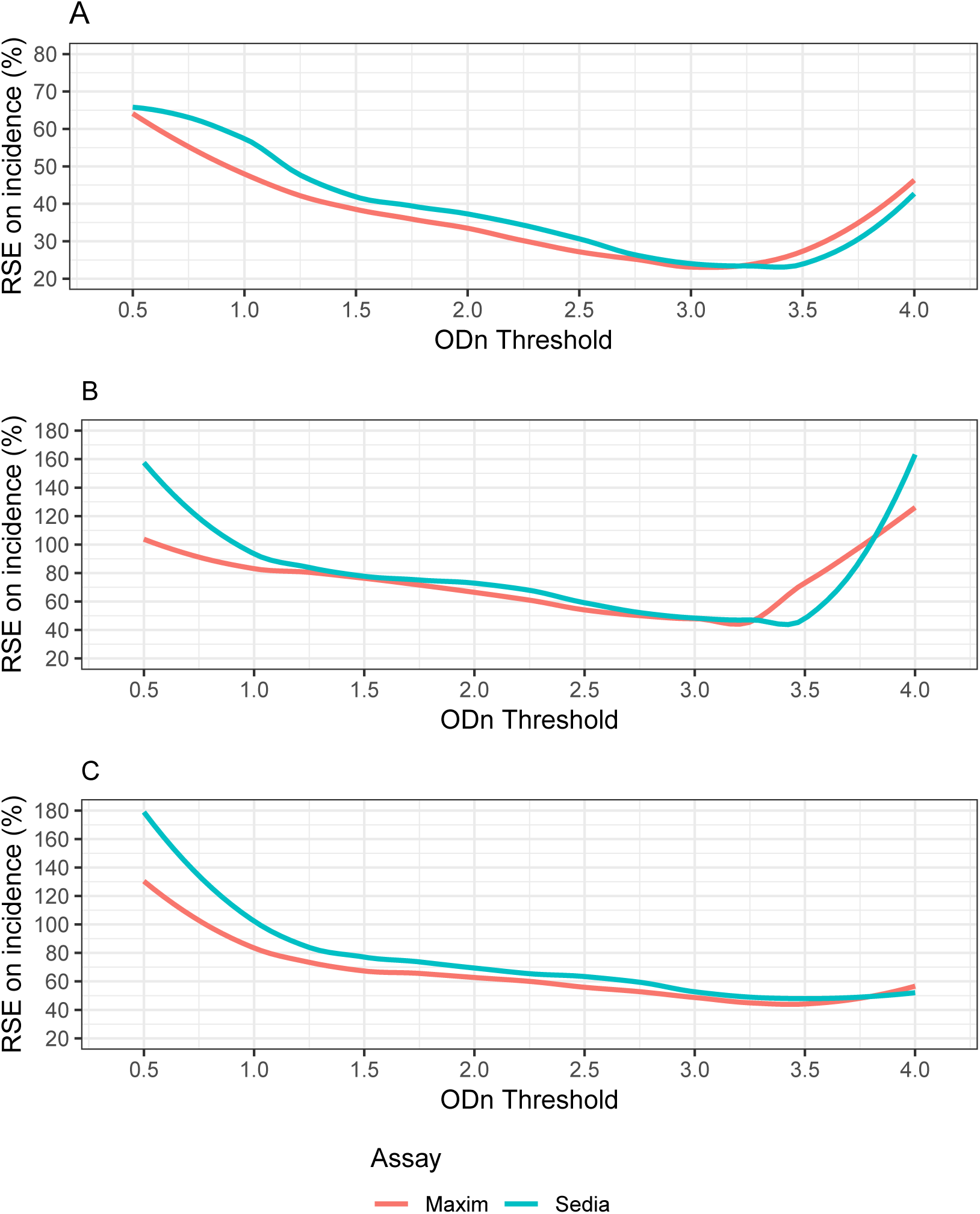
Relative standard error (RSE) of incidence estimate against ODn threshold in three demonstrative surveillance scenarios (assuming imperfect ARV exposure testing) Assuming imperfect ARV exposure testing which reduces the FRR in virally unsuppressed treated individuals to 10% of FRR in suppressed individuals (where no viral load threshold applied). **A.** Scenario similar to South African epidemic. **B.** Scenario similar to Kenyan epidemic. **C.** Scenario similar to North American key population epidemic.

